# HSP70 chaperones IL-33 in chronic airway disease

**DOI:** 10.1101/2025.01.30.635799

**Authors:** Omar A. Osorio, Colin E. Kluender, Heather E. Raphael, Ghandi F. Hassan, Lucy S. Cohen, Deborah F. Steinberg, Ella Katz-Kiriakos, Morgan D. Payne, Ethan M. Luo, Jamie L. Hicks, Derek E. Byers, Jennifer Alexander-Brett

## Abstract

IL-33 is a key driver of type 2 inflammation relevant to airway epithelial biology. However, the mechanisms for IL-33 secretion and regulation in the context of chronic airway disease is poorly understood. Here we report a disease-associated isoform IL-33^Δ34^ that escapes nuclear sequestration to be tonically secreted by recruitment to unconventional protein secretory pathways by HSP70. IL-33^Δ34^ interacts with HSP70 within cells to be targeted to secretory organelles through coordinated binding to phosphatidylserine (PS). Once secreted, this HSP70/cytokine complex stabilizes IL-33^Δ34^ by inhibiting oxidation/degradation thereby enhancing IL-33^Δ34^-receptor binding and activity. We find evidence for IL-33, HSP70 and chaperonin TCP-1 are dysregulated in human chronic airway disease. This phenomenon is reflected in differential proteomics of diseased bronchial wash extracellular vesicles. This study confirms proteostasis intermediates, chiefly HSP70, as a chaperone for non-canonical IL-33^Δ34^ secretion and activity that may be amenable for therapeutic targeting in airway diseases such as asthma and COPD.

**GRAPHICAL ABSTRACT:** 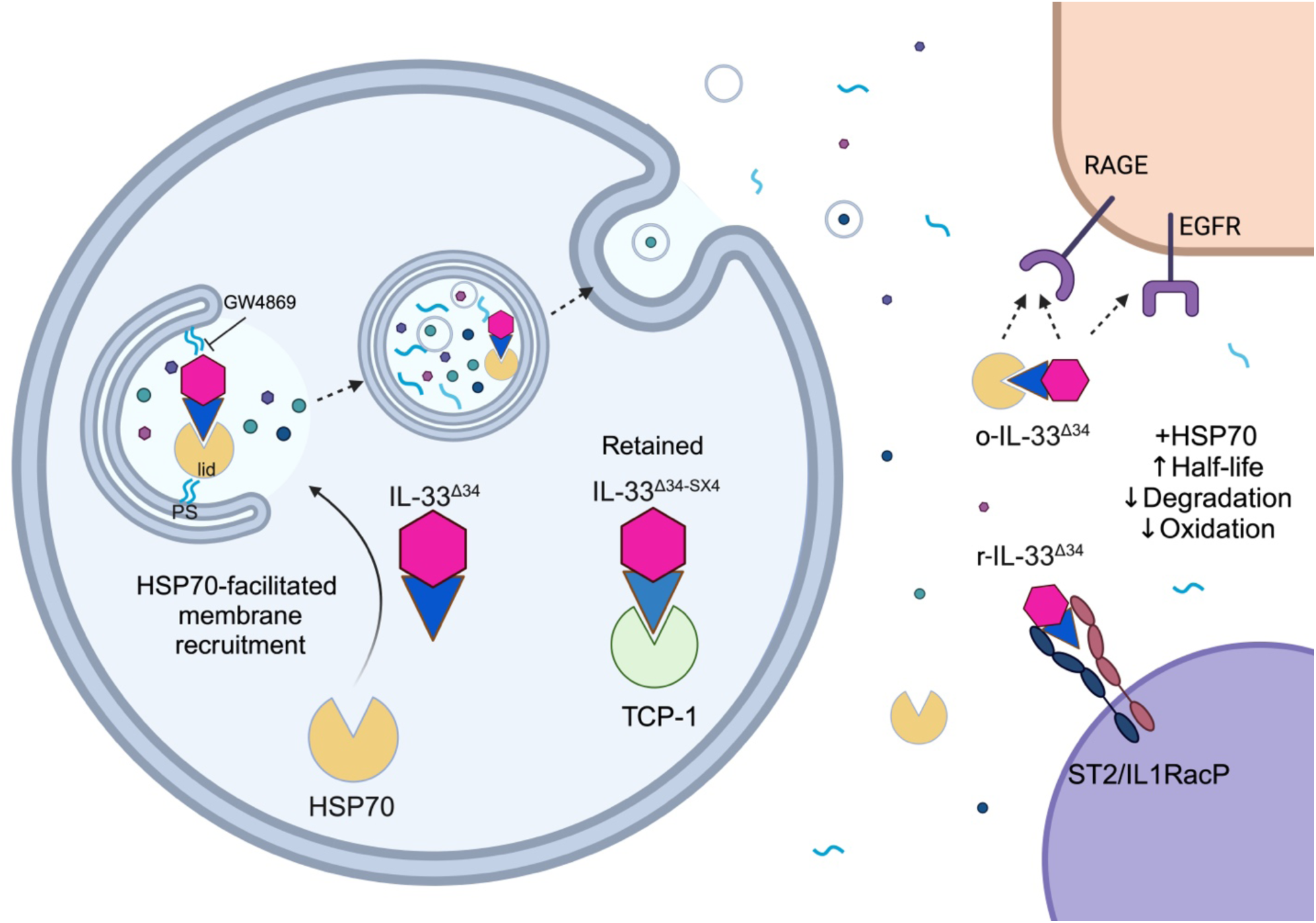

## INTRODUCTION

IL-33 is an epithelial-derived cytokine (1) that can polarize inflammatory responses toward a type-2 immune phenotype (2, 3). Support for IL-33 as central mediator of human airway disease is derived from genome-wide association studies (4–6), and analysis of specimens from human asthma (7, 8) and chronic obstructive pulmonary disease (COPD) patients (9–12). Several animal models of airway disease have proven useful to study the role of IL-33 expression and release as a trigger for airway disease in response to virus infection (9) and allergen challenge (12–14).

The mechanism of IL-33 secretion has been investigated under a variety of conditions, including in response to danger signals, under conditions of cell necrosis or infection models (reviewed in (15)). We previously analyzed the mechanism of respiratory epithelial IL-33 secretion in chronic airway disease, focusing on airway basal cells as the primary cellular source of this cytokine in human lungs. We observed tonic secretion of disease-associated spliced IL-33 isoforms from basal cells, which was driven by the neutral sphingomyelinase-2 (nSMase2)-dependent multivesicular endosome (MVE) pathway. We also found that IL-33 could be co-secreted non-covalently bound to small extracellular vesicles (sEVs) (∼100-150 nm diameter) that exhibited features consistent with exosomes of endolysosomal origin (12, 16). We found nSMase2 to be increased in COPD-derived specimens and demonstrated attenuated IL-33 driven type-2 inflammation under conditions of nSMase2 blockade in an Alternaria model.

HSP70 (HSPA1A (17)) is an inducible member of the Heat Shock Protein (HSP) family that broadly function as chaperones for nascent and misfolded proteins, translocation of polypeptides across organelle membranes, and the targeting of proteins for degradation by the ubiquitin-proteasome system or via chaperone mediated autophagy (CMA) (18–20). HSP70 expression is rapidly upregulated in response to cellular stress to overcome environmental damage and pathophysiology (21, 22). In the context of airway disease, increased expression levels of HSP70 have been observed in COPD lung tissue (23) matched by higher extracellular HSP70 protein measured in sputum and serum or plasma of patients with asthma COPD and asthma-COPD overlap (ACO) (24, 25). HSPs are also extensively involved in unconventional protein secretion in leaderless proteins, like the unconventional secretion of IL-1 family member IL-1Δ (26–28). Herein, we demonstrate that HSP70 interacts with IL-33^Δ34^ to facilitate its recruitment to unconventional protein secretion pathways in airway epithelial cells. Once outside of the cell HSP70 appears to stabilize IL-33 by decreasing oxidation/degradation resulting in enhanced receptor signaling. The involvement of heat shock pathways in airway disease is supported by altered expression and localization of IL-33 and heat shock proteins in lung tissue and extracellular vesicle proteomic profiles.

## RESULTS

### IL-33^Δ34^ Interacts with HSP70

Previous studies have demonstrated the alternative splicing of IL-33 in diseased human airway cells and lung tissue (12, 29). We expressed these variants in mammalian Expi293 cells with dual FLAG and 6xHis tags and performed tandem affinity purification, which revealed a co-eluting band of high molecular weight for all IL-33 isoforms (Fig 1A). We identified this co-eluting band as containing HSP70 and HSC70 by mass spectrometry analysis (Fig 1B), with ratio depending on cell type analyzed. We validated this by co-immunoprecipitation using purified recombinant biotinylated IL-33^Δ34^ and HSP70 expressed in E. coli (Fig 1C). Interestingly, there is reduced interaction when an oxidation resistant form of IL-33^Δ34^ (IL-33^Δ34SX4^) (30) is used as the bait. We demonstrated close intracellular interactions (∼10-15 nm labeling radius (31)) between these two proteins by expressing an IL-33^Δ34^ miniTurboID fusion protein proximity ligation assay (32) (Fig 1D). This appears to occur in a cell-type specific manner as the same fusion construct resulted in minimal HSP70 biotinylation in a U937 human monocytic cell line. Based on our observations for the oxidation resistant form of IL-33, we performed comparative proximity ligation analysis, which revealed differential labeling for several heat shock and proteostasis intermediates (examples include HSP70/HSPA1A, HSC70/HSPA8, FLNA/B, ANXA1/2, and TCP1 subunits CCT2, 3, 4,5, 7 and 8).

**Figure 1.**
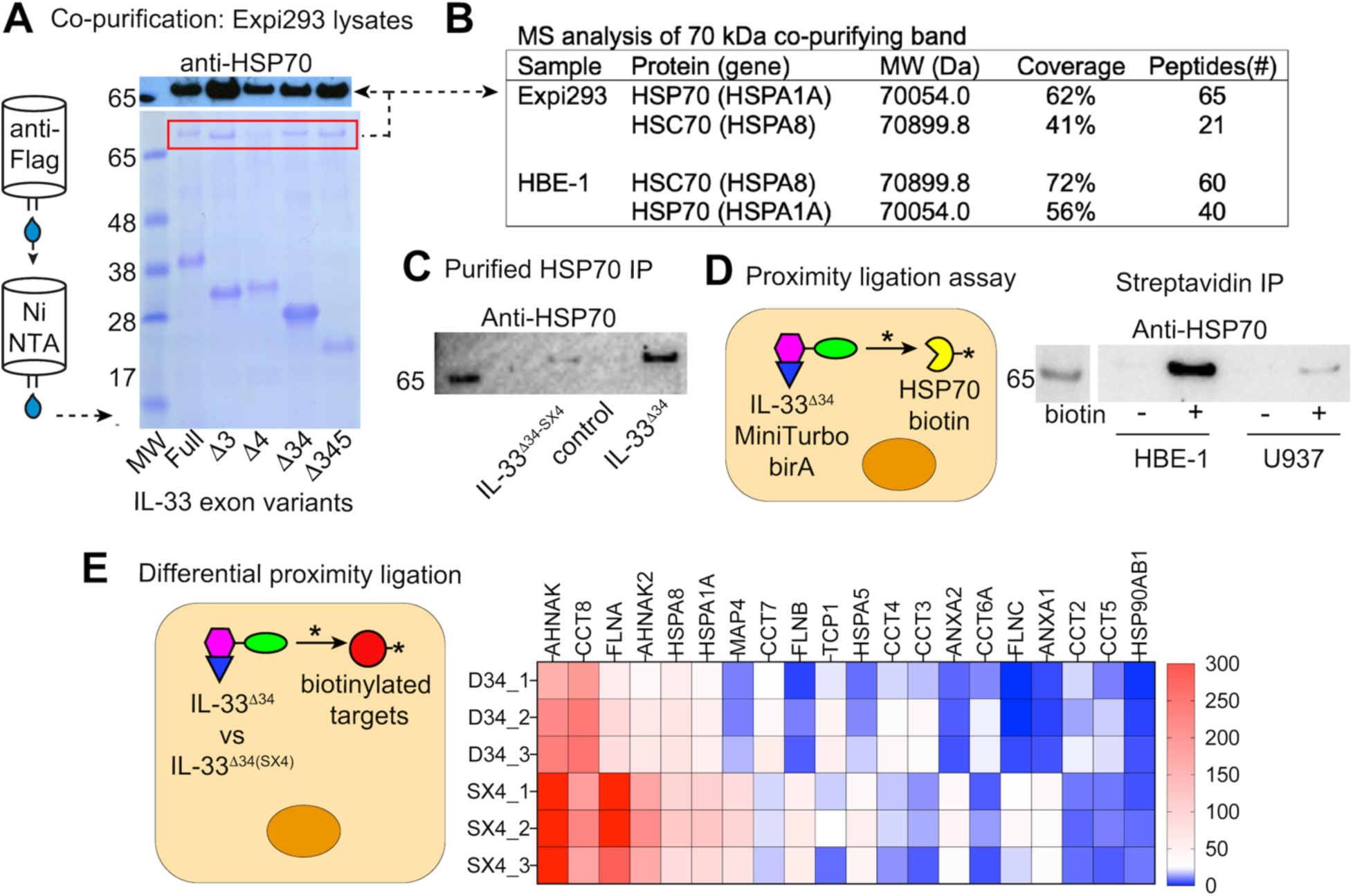
HSP70 interacts with IL-33 in human airway epithelial cells. A) SDS-PAGE gel demonstrating co-purification of a high molecular weight band (>65 kDa, MW markers as indicated) that co-purified with all IL-33 isoforms on tandem anti-FLAG, NiNTA affinity purification. B) Mass Spectrometry analysis of contaminating band isolated from Expi293 expression cells and HBE-1 airway cell line identified as HSP70 (and HSC70) with coverage and relative # peptides reported. C) Recombinant purified human IL-33^Δ34^ and HSP70 interact by Co-Immunoprecipitation of biotinylated IL-33 protein compared to oxidation resistant Cys->Ser x4 mutant IL-33^Δ34SX4^. D) Intracellular interaction of these proteins verified based on overexpression of miniTurboID-IL-33^Δ34^ proximity ligase fusion protein with robust biotinylation of HSP70 in HBE-1 cells but not in U937 cells. E) Differential proximity ligation comparison for IL-33^Δ34^ and IL-33^Δ34SX4^ miniTurboID fusions in HBE-1 cells. Heatmap of top 20 most abundant proteins and additional heat shock intermediates of relevance. Peptide number indicated by color mapping blue (0), white (25), red (300), replicates 1-3 as shown.

### HSP70 Facilitates IL-33^Δ34^ Secretion

We previously reported that IL-33^Δ34^ is tonically secreted from primary airway basal cells and cell lines, which is blocked by GW4869 (12). We therefore utilized this spontaneous secretion cytokine variant to test IL-33 secretion efficiency by disrupting its interaction with HSP70. We first demonstrated reduced secretion into the cell supernatant for the oxidation resistant IL-33^Δ34-SX4^ form in HBE-1 cells compared to cysteine containing IL-33^Δ34^ (Fig 2A, Fig S1E). We also measured secretion in the context of the immune cell line U937 that also exhibited reduced HSP70 labeling by proximity ligation and noted similarly the immune U937 cells did not secrete IL-33^Δ34^ into cell supernatant (Fig 2B).

**Figure 2.**
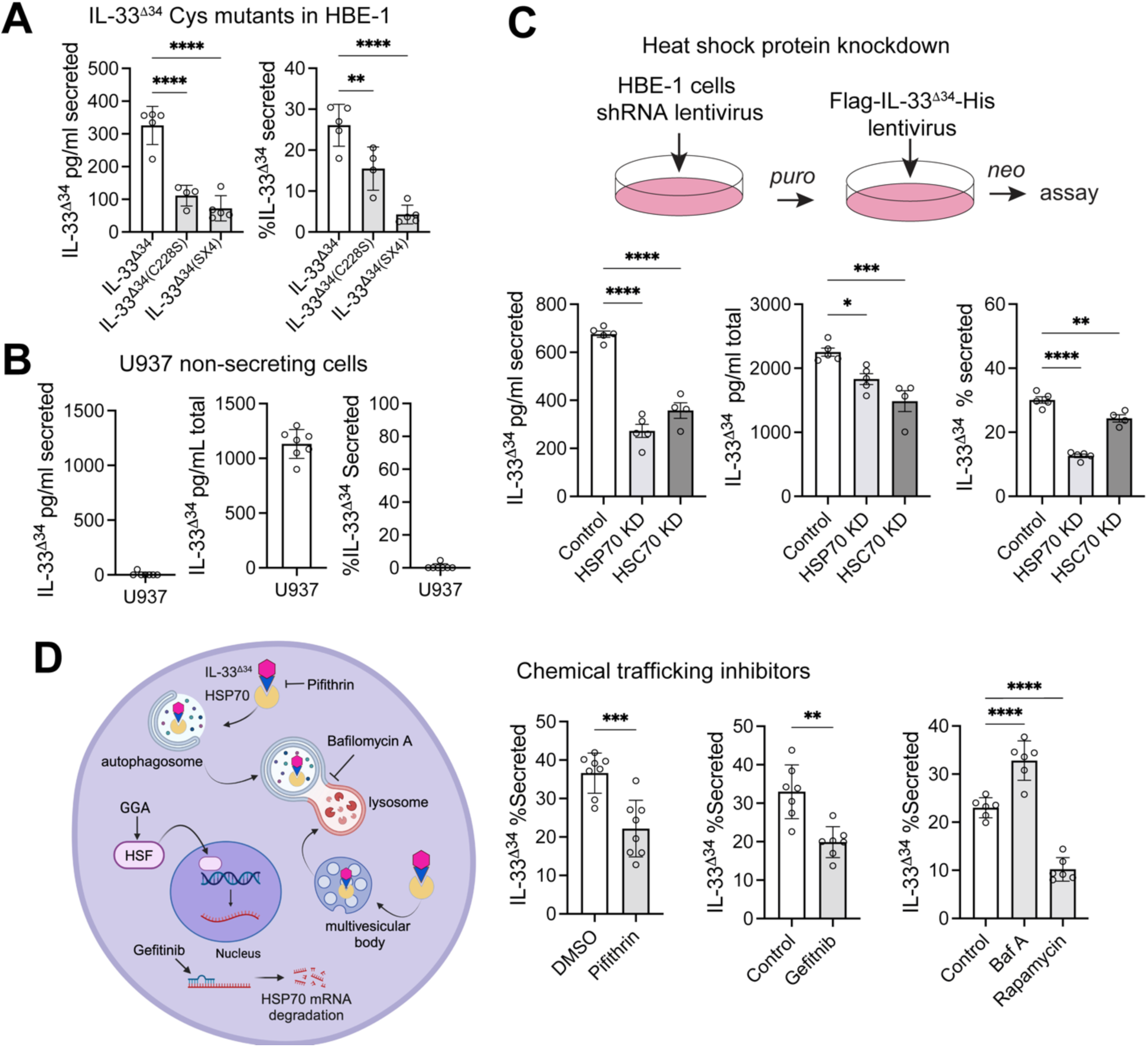
HSP70 facilitates IL-33 secretion. A) HBE-1 cell secretion by ELISA for lentiviral-expressed IL-33^Δ34^, IL-33^Δ34SX4^ and IL-33^Δ34C228S^ proteins represented as pg/ml secreted into supernatant and secretion efficiency normalized for total protein expression level. B) ELISA IL-33^Δ34^ secretion assay in U937 cells that demonstrate no detectable secretion despite robust intracellular expression in the ng/ml range. C) HBE-1 airway cells were co-transduced with lentiviruses expressing shRNA targeting HSP70 or HSC70 and dual selected for ELISA secretion assay. D) Schematic of HSP70 and vesicular trafficking inhibitors. ELISA secretion assay performed with HSP70 inhibitors: Pifithrin-μ (10 μM), Gefitnib (1 μM), autophagosome acidification inhibitor Bafilomycin A (Baf A) and autophagy activator Rapamycin (100 nM). Inhibitors were pre-incubated on cells per Methods prior to secretion assay. Data points displayed with mean ± SEM. *P*-value limits: \**P* < 0.05, \*\**P* < 0.01, \*\*\**P* < 0.001, \*\*\**P* < 0.0001.

To determine if HSP70 interaction is required for IL-33^Δ34^ secretion, we measured secretion in the context of HSP70 and HSC70 knockdown in HBE-1 cells. We were able to robustly knockdown HSP70 mRNA (Fig S1A) and observed a modest decrease of IL-33^Δ34^ secretion into the supernatant when normalized to total IL-33^Δ34^ (Fig 2C). This modest effect is attributed in part to very high baseline expression levels of HSP70 and HSC70 in our cell lines. We therefore supplemented this with an orthogonal chemical blockade approach, disrupting the HSP70 axis using Pifithrin-μ and Gefitinib, inhibitors of HSP70 substrate binding and expression respectively (33, 34). Both reduced the secretion efficiency of IL-33^Δ34^ in our assay (Fig 2D, Fig S1B).

HSP70 has also been implicated in non-classical secretion mechanisms such as secretory autophagy or the extracellular vesicle (EV) pathways through chaperone-mediated autophagy (CMA) or endosomal microautophagy (35, 36). Using inhibitors of these systems, we saw a significant increase in IL-33^Δ34^ release with bafilomycin A (Baf A) treatment, a chemical which inhibits autophagy and increases extracellular vesicle release (37). We observed a corresponding decrease in IL-33^Δ34^ secretion with rapamycin, which promotes autophagy and decreases extracellular vesicle release (38) (Fig 2D, Fig S1C).

### IL-33^Δ34^ can bind Phosphatidylserine

Neutral sphingomyelinase-2 (nSmase2) is activated by Phosphatidylserine (PS) binding and a recent structural study demonstrated that GW4869 is a lipid competitive inhibitor (39). Given our previously observed blockade of IL-33^Δ34^ using the nSMase2 inhibitor GW4869, we employed a lipid binding ELISA assay to test whether the compound could also influence direct cytokine interactions with lipids (schematic Fig 3A). We found IL-33^Δ34^ readily bound to PS and that this was significantly abrogated in the presence of GW4869 (Fig 3B). We also observed that the non-secreted IL-33^Δ34SX4^, has significantly reduced ability to bind PS (Fig S2A) compared to IL-33^Δ34^. We tested whether oxidation could impact lipid binding by IL-33^Δ34^ (o-IL-33^Δ34^), in which the surfaced exposed cysteines have undergone disulfide bridge formation and consequent confirmational change that inactivates the cytokine for IL-1RL1 binding (30). We found oxidation reduced IL-33^Δ34^ affinity for PS (Fig 3B) but the addition of HSP70 was able to rescue PS binding by o-IL-33^Δ34^. HSP70 and HSC70 have both been shown to bind PS (40, 41) but unlike IL-33^Δ34^, GW4869 does not appear to block the release of HSP70 (Fig 3C) which may indicate specificity of GW4869 for specific lipid-binding proteins or inadequate dosing given endogenous HSP70 protein is expressed at 2 orders of magnitude higher levels than overexpressed IL-33^Δ34^ within HBE-1 cells, with similar high extracellular levels in cellular assay (Fig S2D).

**Figure 3.**
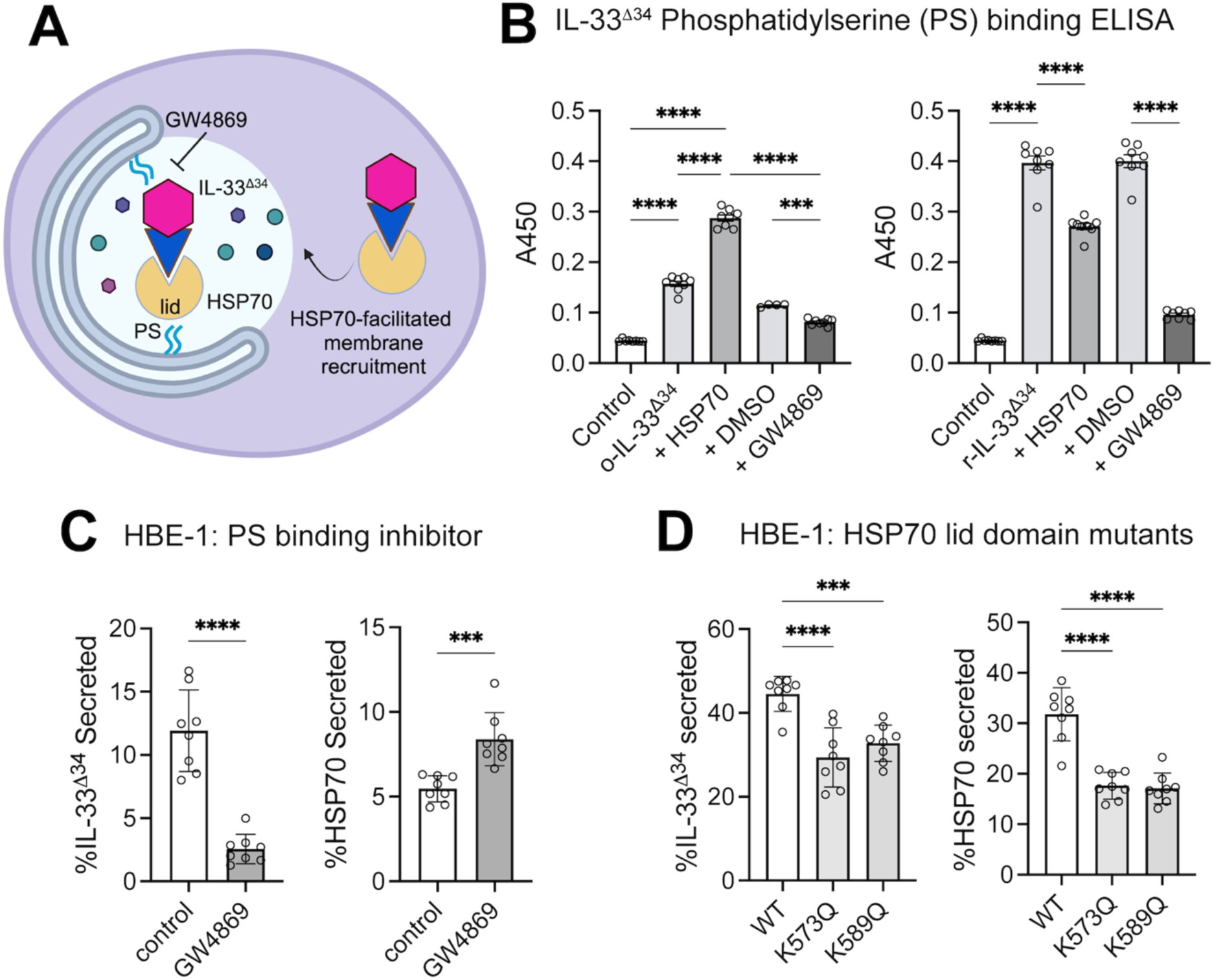
IL-33 Interacts directly with phospholipids. A) Schematic of model for phosphatidylserine (PS) cooperative recruitment of IL-33^Δ34^ and HSP70 to a secretory vesicular compartment. B) PS binding ELISA performed using reduced (r-IL-33^Δ34^) and oxidized (o-IL-33^Δ34^) cytokine (100 ng/ml) +/- HSP70 (300 ng/ml) or GW4869 (20 μM) or DMSO vehicle control. C) Secretion ELISA data for IL-33^Δ34^ and HSP70 under conditions of GW4869 treatment. D) Secretion ELISA data for IL-33^Δ34^ under conditions of lentiviral overexpressed HSP70 LID domain mutants K573Q and K583Q. Data points displayed with mean ± SEM. *P*-value limits: \**P* < 0.05, \*\**P* < 0.01, \*\*\**P* < 0.001, \*\*\**P* < 0.0001.

Prior structural analyses have identified key residues in the HSC70 LID domain (41) responsible for binding to PS. We therefore overexpressed two PS binding-deficient mutants K573Q and K589Q in the context of HSP70 in HBE-1 cells and observed a reduction in both IL-33^Δ34^ and HSP70 in the conditioned media compared to WT HSP70 (Fig 3D, Fig S2E).

### HSP70 differentially modulates IL-33 binding and signaling through receptors

Since HSP70 is also secreted from cells (18), we next asked whether this association would influence IL-33^Δ34^ biology outside of the cell. We observed that HSP70 did not affect reduced (r-IL-33^Δ34^) binding to IL-1RL1/ST2 (Fig 4A) by ELISA nor did it impact IL-1RL1/IL1RAP signaling in a HEK-Blue reporter cell line (Fig 4B). However, oxidized IL-33 has been attributed to binding and signaling through a novel receptor complex of RAGE and EGFR (42), and HSP70 is a known RAGE ligand (43). We therefore tested whether HSP70 impacted o-IL-33^Δ34^ binding to RAGE binding and found that indeed HSP70 reduced o-IL-33^Δ34^ interaction with RAGE by ELISA (Fig 4A) and reduced RAGE signaling. As anticipated, HSP70 alone exhibited the most robust signaling in the A549 RAGE signaling reporter line but had no effect on signaling in the HEK-Blue IL-1RL1 reporter line. Notably, oxidized-IL-33^Δ34^ was unable to bind IL-1RL1, but addition of HSP70 during oxidation step did partially rescue IL-1RL1 signaling (Fig 4B). To complement ELISA-based receptor binding, we tested IL-33^Δ34^ binding to cell surface receptors on A549 and HEK-Blue cell lines. We observed an increase in IL-33 fluorescence signal in the presence of HSP70 for both cell lines, which was relatively decreased in the presence of soluble RAGE or sST2 (Fig 4C). This effect was more pronounced for RAGE likely due to relative binding affinity of overexpressed IL-1RL1-IL1RAP complex on HEK-Blue reporter cells. The significant influence of HSP70 on IL-33^Δ34^ signaling, and cell-surface binding led us to test if HSP70 helps stabilize IL-33^Δ34^ in the extracellular environment. In fact, co-incubating HSP70 with IL-33^Δ34^ in serum-free media protects IL-33^Δ34^ from both oxidation (seen as a duplicate band just below the expected 28kDa IL-33^Δ34^ band) and degradation (seen as increased laddering and loss of signal) (Fig 4D).

**Figure 4.**
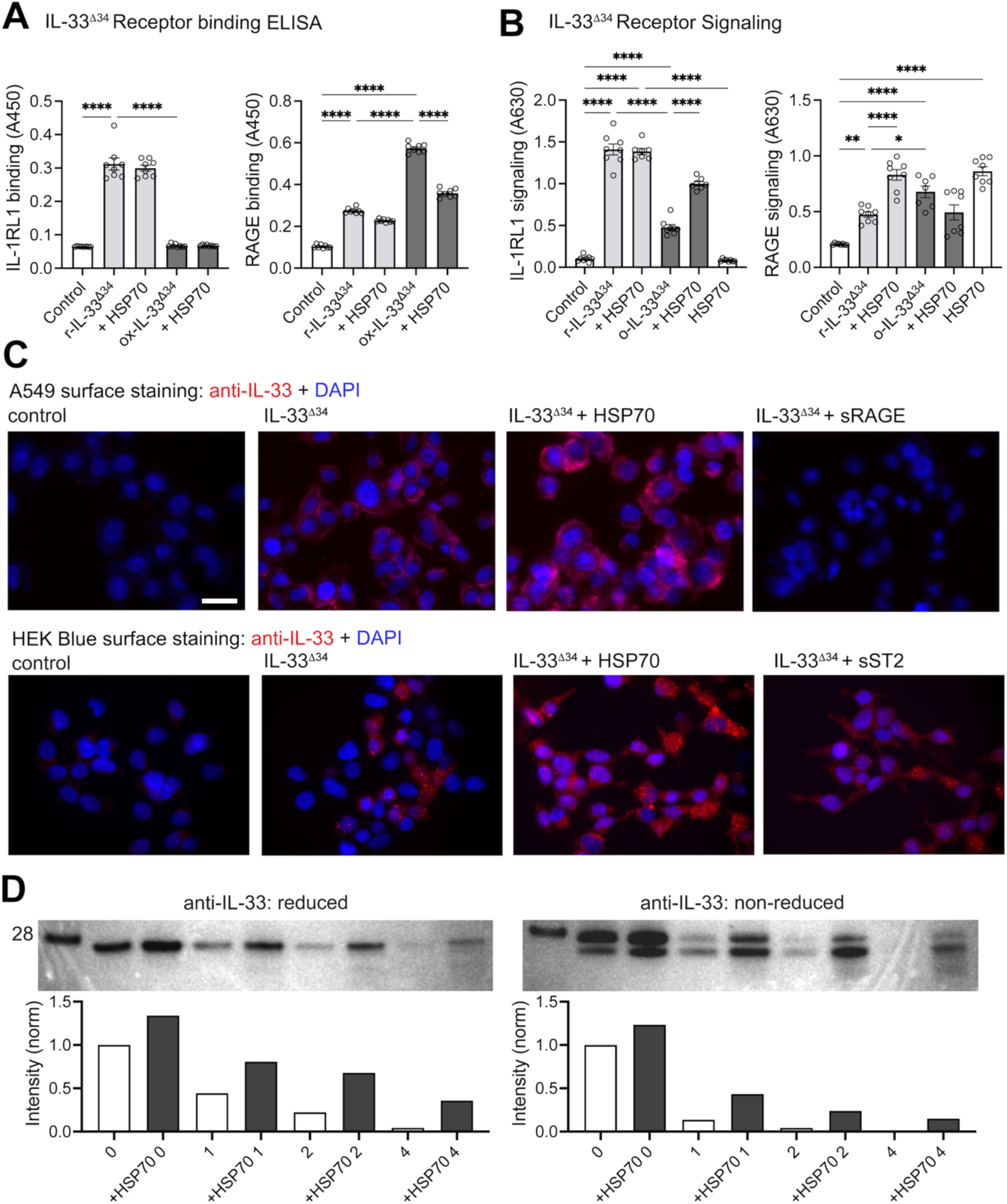
HSP70 preserves oxidized IL-33 receptor signaling. A) Receptor-binding ELISA assays for IL-1RL1 and RAGE binding to reduced (r-IL-33^Δ34^) and oxidized (o-IL-33^Δ34^) cytokine (100 ng/ml) +/- HSP70 (300 ng/ml) and with HSP70 alone (known ligand for RAGE, not IL-1RL1). B) Cellular signaling assays performed under same conditions as in A) using HEK293T (HEK Blue, overexpressed IL-1RL1-IL1RAP complex and AP-1/NF-κB reporter) or A549 Dual (endogenous RAGE, NF-κB reporter) cell lines with development of secreted alkaline phosphatase (SEAP) as readout. C) Immunofluorescence staining of A549 Dual and HEK Blue cells demonstrating IL-33^Δ34^ protein binding (red) to the cell surface +/- HSP70 or soluble forms of corresponding receptors RAGE and IL-1RL1. DAPI nuclear counterstain (blue). Imaging performed at 60X magnification, scale bar 20 μm. D) IL-33^Δ34^ was diluted in media (1 μg/ml) +/- HSP70 (10 μg/ml) and aliquots taken at 0, 1, 2 and 4 hr timepoints for reduced and non-reduced immunoblots, quantification of band intensity below gel images with corresponding lane conditions. Data points displayed with mean ± SEM. *P*-value limits: \**P* < 0.05, \*\**P* < 0.01, \*\*\**P* < 0.001, \*\*\**P* < 0.0001.

### Heat shock transcripts are modulated in airway disease

We have previously shown that *IL33^Full^* and *IL33^Δ34^* mRNA expression is upregulated in COPD lung tissue (12, 44). Here we analyzed a cohort of specimens across a spectrum of airway disease severity. We have collected and characterized a large cohort of airway specimens including nonsmokers, smokers, mild-moderate COPD and severe COPD, as well as severe asthma (Table S1). We measured expression levels of several heat shock and proteostasis intermediates in this cohort by qPCR (Fig 5A and Fig S3A). We found, expectedly, that *IL33*, *SPMD3* (nSmase2) and *MUC5AC* were increased with smoking and progression of airway disease and correlated with increased *IL33* expression (Fig S3B). Interestingly, *SMPD3* expression was prominently and specifically increased in severe COPD and not modulated in the (smaller) asthma cohort. And while we did not observe statistically significant changes in *HSPA1A* or *HSPA8* transcript levels, other stress induced intermediates including *HSP90AA1* and the heat shock cochaperone *STIP1* (45) were upregulated in severe COPD (Fig 5A). Other vesicular trafficking intermediates tested were not differentially expressed, although the proteostasis moderator p62/*SQSTM1* exhibited a trend toward decreased expression in severe disease (Fig S3A).

**Figure 5.**
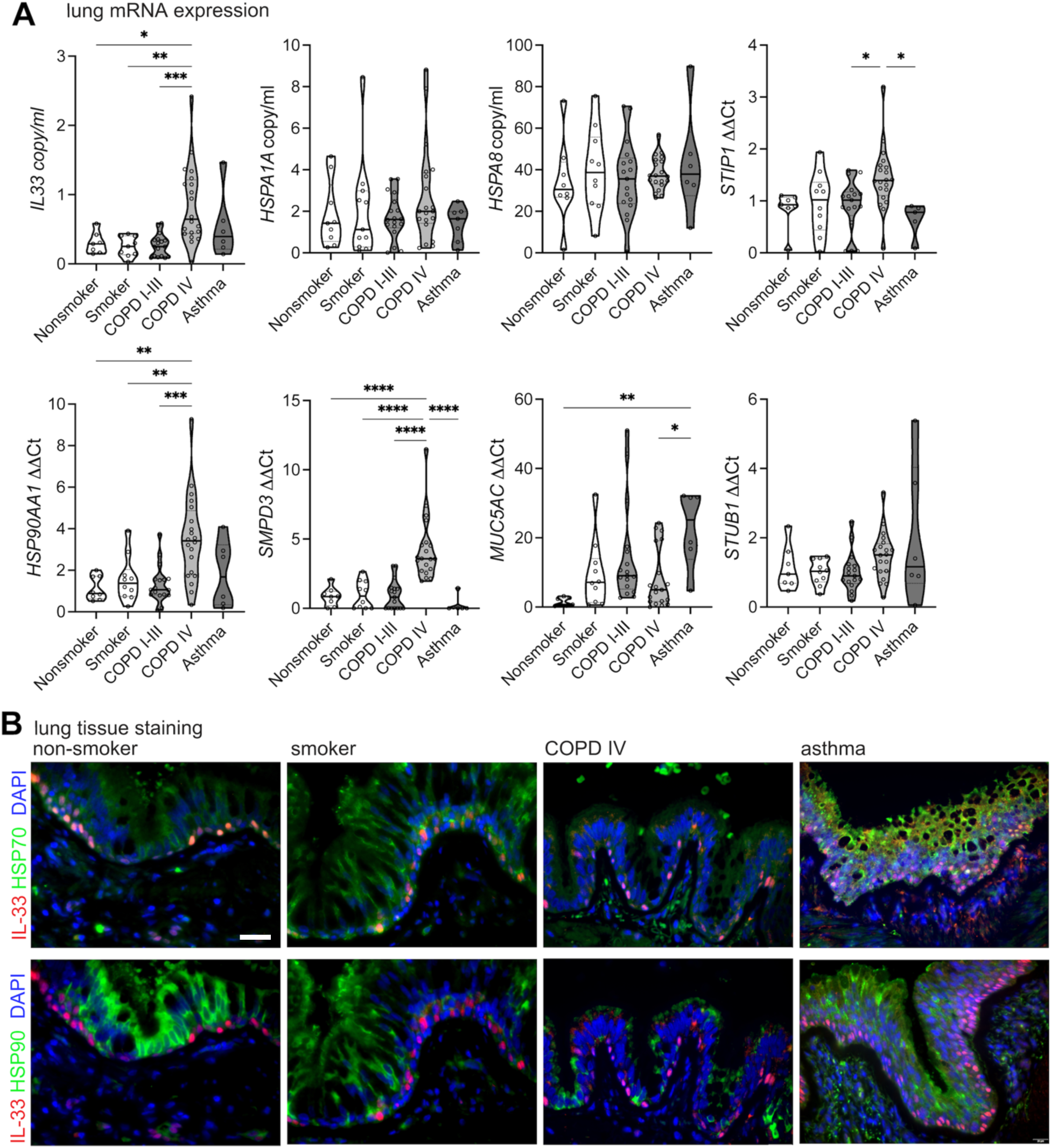
Heat shock intermediates and IL-33 expression in airway disease tissue. A) Human lung tissue specimens were stratified based on GOLD classification and smoking status for non-diseased specimens (see Table S1). Transcript expression was measured by gene-specific qPCR for nonsmokers (n=7), smokers (n=11) COPD I-III (n=17), COPD IV (n=20) and severe asthma (n=6) lung tissue specimens represented as copy/ml based on plasmid standard or fold-change by ΔΔCt method as shown, normalized to GAPDH. B) Immunofluorescence staining for representative human lung tissue specimens across COPD disease severity and for asthma specimens. Staining for IL-33 (red) and HSP70 or HSP90 (green) as indicated; DAPI counterstain (blue), imaging performed at 40X magnification, scale bar 50 μm. Data points displayed with mean ± SEM. *P*-value limits: \**P* < 0.05, \*\**P* < 0.01, \*\*\**P* < 0.001, \*\*\**P* < 0.0001.

### Heat shock proteins are altered in airway disease

To complement transcript level expression data, we also investigated protein expression patterns for heat shock intermediates in airway disease tissue. We found HSP70 showed diffuse localization throughout the epithelium but was present and relatively concentrated within IL-33 expressing basal cells (Fig 5B). In contrast, HSP90 protein exhibited intense immunostaining in differentiated airway cells, with relative exclusion from IL-33 expressing airway basal cells. Comparison across disease states demonstrates relative decrease in HSP70 staining in severe COPD specimens, with diffuse, patchy staining in areas of airway basal hyperplasia in asthma. By comparison, HSP90 protein staining was also decreased in severe COPD airways, with a relative paucity of staining observed in asthma as well, which appears to reflect the abundant mucus metaplasia in both severe COPD and asthma. These results indicate that intracellular protein levels of heat shock intermediates appear to be uncoupled from transcript expression in airway disease and are impacted by COPD disease severity and airway disease phenotype.

We also measured total protein concentrations of IL-33, HSP70 and HSP90 in diseased tissue lysates and bronchial wash fluid. As anticipated, IL-33 protein levels trended upward with disease severity, while HSP70 amounts decreased in an inverse manner (Fig 6A). Despite decreases in lysate HSP70 levels, bronchial wash (BW) HSP70 concentrations remained relatively constant (Fig 6B) though the ratio of BW to total lung HSP70 as an indirect measure of extracellular vs intracellular fractions was significantly increased in severe COPD. A similar result was observed for HSP70 concentrations measured in BAL from the SPIROMICS cohort, though the absolute concentration of protein was 10-fold lower, likely a dilutional effect (Fig 6C). Notably, severe COPD was under-represented in the SPIROMICS cohort. For HSP90, lung tissue and BW specimens exhibited a very broad range of concentrations with significant decreases in BW fluid for all disease states compared to nonsmokers (Fig S3C). A similar non-significant decreasing trend was observed in SPIROMICS BAL specimens. These trends likely reflect the process of mucus metaplasia as a function of disease severity, as observed for tissue immunostaining. Additionally, this variability observed may be related to decreased detection of HSP90 due to incorporation into small EVs (46).

**Figure 6.**
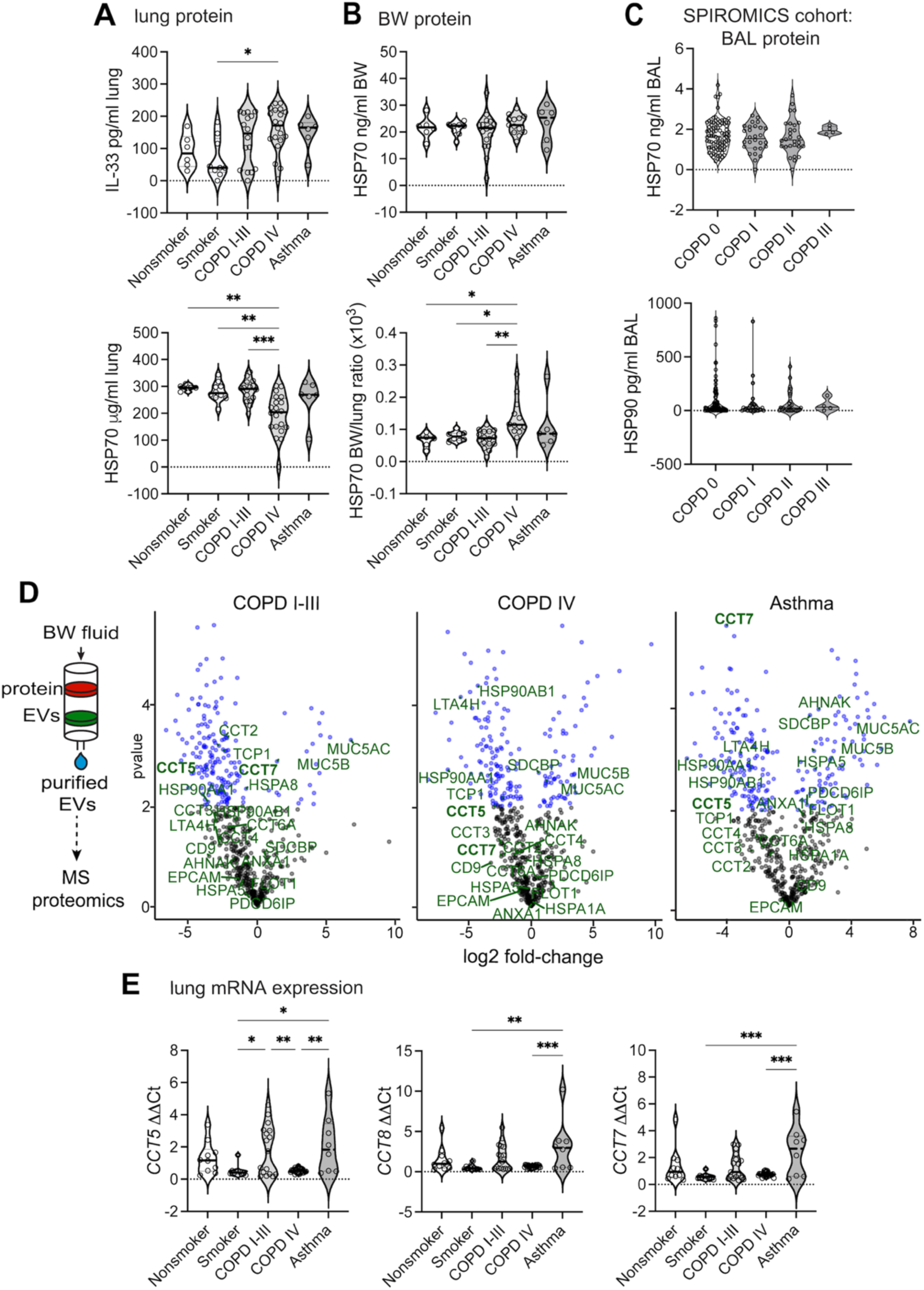
Heat shock intermediates and IL-33 are present in bronchial lavage fluid and EVs contents are altered in disease. A and B) Protein levels for IL-33 and HSP70 in lung tissue and lavage fluid measured by ELISA. C) HSP70 and HSP90 protein levels measured by ELISA in SPIROMICS BAL specimens. D) Schematic for EV isolation by size exclusion chromatography from bronchial wash (BW) and volcano plots for differential protein expression in COPD IV, COPD I-III and asthma specimens compared to nonsmoker control. Proteins annotated are associated with heat shock or proteostasis systems and were biotinylated by IL-33 in miniTurboID experiment (Fig 1E). Bronchial wash EV proteome demonstrated for HSP70, HSC70 and HSP90. E) Transcript expression of molecular chaperonin TCP1 subunits *CCT5*, *CCT7* and *CCT8* in lung tissue displayed as fold-change by ΔΔCt method with normalization to *GAPDH*. Data points displayed with mean ± SEM. *P*-value limits: \**P* < 0.05, \*\**P* < 0.01, \*\*\**P* < 0.001, \*\*\**P* < 0.0001.

### Differential EV-mediated secretion of heat shock proteins in airway disease

The common link between GW4869, increased nSmase2 activity and heat shock proteins is well established in the field of extracellular vesicle biology with a prominent link to EVs of endolysosomal origin (46). We have previously shown that IL-33^Δ34^ can interact with purified EVs isolated from lung lavage fluid (12) and therefore isolated bronchial wash EVs from specimens across the spectrum of COPD disease severity to examine the proteome for comparison to our transcript and protein expression data. We isolated EVs as per our established protocols (12) and validated our preparation with the ExoView SP-IRIS imaging platform (Fig S3D). We analyzed differential proteomics in EVs isolated from BW fluid from nonsmoker, COPD I-III, COPD IV and asthma specimens (Fig 6D). As expected, we observed differentially increased MUC5AC and MUC5B mucins in COPD. While HSP70 and HSC70 were not differentially present, HSP90 was decreased in correlation with down-trending protein levels in BW and BAL fluid as well as diminished immunostaining in airway disease. Also of relevance, multiple intermediates of leukotriene metabolism (LT4AH, ALOX15) were present in BW EVs. Of significance, we found multiple proteins from the TCP1 proteostasis-degradataion complex (CCT2, CCT4, CCT5, CCT7, CCT8) were decreased in airway disease BW EVs as well as epithelial markers (EPCAM). Comparison of these findings to miniTurboID proximity assays in Fig 1E suggests a potential role for TCP1 complex proteins in IL-33 cellular retention. Accordingly, expression levels for TCP1 intermediates with differential EV incorporation were also modulated in severe COPD on the lung tissue level (Fig 6E), suggesting this could be another component of cellular proteostasis machinery that is dysregulated in airway disease.

## DISCUSSION

Heat shock proteins and proteostasis pathways have been long linked to chronic airway disease pathogenesis (23–25, 47). Elevated levels of HSP70 have been reported in plasma and sputum of COPD, asthma and asthma-COPD overlap (ACO) patients and in the BAL of induced airway inflammation in mouse models (47). This natural association stems from the role of these proteins in response to cellular stress, toxin-mediated tissue damage and hypoxia which are common in the diseased airway tissue environment. It has long been thought that HSP70 has an anti-apoptotic effect and as part of the cellular stress response functions to bring cells back into homeostasis (48). Due to its vital intracellular role and lack of signal peptide led to extracellular HSP70 being categorized as a DAMP that induced a pro-inflammatory response in what was called the Chaperone Balance Hypothesis (49). Succinctly, this hypothesis states that the inflammatory effect of HSP70 was determined by the ratio of extracellular to intracellular HSP70 concentrations and that the more the HSP70 pool moved into the extracellular compartment, the more pro-inflammatory effect it exerted and the shift in compartmental concentrations is evident in diseases like type 2 diabetes. When it comes to airway inflammation models, both pro- and anti-inflammatory effects of exogenously administered extracellular HSP70 have been reported, meaning that we still do not know what function/effect extracellular HSP70 has on the airway in the context of chronic airway disease.

In this study, we link HSP70 to recruitment of IL-33 for non-canonical secretion from airway cells and potential stabilization outside of the cell. Additional work will be necessary to fully characterize the effect of HSP70 on IL-33 oxidation and sensitivity to protease degradation, but our results presented here identify a previously unrecognized function for heat shock proteins in facilitating the secretion and function of a pro-inflammatory cytokine. We have further demonstrated that this function requires the presence of cysteine residues on IL-33, suggesting a potential for covalent HSP70 adducts to form with clients as part of this stabilization function. Such an adduct may hold IL-33 in an active conformation in an oxidating environment to maintain potency and promote airway disease. This may be a clue as to how HSP70 binds IL-33^Δ34^ at the molecular level, given that IL-33 does not contain the canonical KFERQ binding motif used by HSPs to bind clients (50).

We have also identified through a combination of proximity ligation assay and bronchial lavage EV proteomics a novel modulatory axis for non-canonical secretion and/or restriction of inflammatory protein secretion through TCP1 complex proteins (51). Our results in distinct cell lines further suggest these pathways may function in a cell type specific manner to regulate directed secretion of EVs, EV cargo, and associated cytokines in respiratory mucosal crosstalk. Future studies will investigate the cell- and tissue-specific functions of the TCP1 complex in airway homeostasis, EV secretion mechanisms and respiratory disease.

## ACKNOWLEDGEMENTS

Special thanks to Jeff Haspel for providing insightful feedback during compilation of this manuscript. We thank the Pulmonary Morphology Core for tissue histology preparation. We thank the Washington University Center Mass Spectrometry Technology Access Center at McDonnel Genome Institute (NIDDK P30 DK020579, NCATS UL1 TR002345, NCI P30 CA091842) for Mass Spectrometry data collection and analysis. Support for this work was provided by NIH/NHLBI (K08HL121168, R01HL52245, R01HL170198 JAB; T32HL007317 to OAO), American Thoracic Society (Early Career Investigator Award, JAB), Burroughs Wellcome Fund (Career Award for Medical Scientist 1014685, JAB), and Doris Duke Foundation (Fund to Retain Clinical Scientists 2015215, JAB).

## AUTHOR CONTRIBUTIONS

O.O., C.K., H.R., G.H., L.C., D.S., M.P., E.L. and J.A.B. designed and/or performed the experiments; O.O. and J.A.B. prepared figures and wrote the manuscript; J.H., E.K. and D.B. contributed to enrollment of human subjects and biobanking efforts.

## METHODS

### Human Lung Samples and Study Design

Clinical samples were obtained from consenting patients at the time of lung transplantation from COPD recipients (n=20) with very severe disease (GOLD Stage IV) during the period from 2016-2025 at Barnes-Jewish Hospital (BJH, St. Louis, MO). Non-diseased (nonsmoker n=7 and smoker n=11), mild-moderate COPD (GOLD I-III) and severe asthma (n=8) specimens were obtained from lungs not suitable for transplantation at BJH (Table S1). There were no pre-determined inclusion or exclusion criteria beyond criteria for lung transplant candidacy. To analyze tissue staining, gene expression and protein levels, lung tissue samples were collected and processed for histopathology, RNA and protein analysis from four different lung zones of each specimen and combined for analysis. Equivalent quantities from the 4 lung areas were pooled for RNA and protein analysis to represent a single sample per specimen. Tissue was homogenized in Trizol (Invitrogen) for RNA extraction or minced and lysed in T-PER (Pierce) supplemented with HALT protease inhibitor (Pierce) for protein analysis. Tissue specimens were fixed in 10% neutral buffered formalin (ThermoFisher) prior to paraffin embedding and sectioning for histopathology analysis. Bronchial wash (BW) fluid was obtained from explanted lungs by instilling 100 ml of PBS into mainstem bronchi and fluid recovered with passive return and gentle suctioning. Bronchial wash fluid was centrifuged at 100x*g* to pellet cells, HALT was added to supernatant prior to storage for further analysis. Bronchoalveolar lavage fluid was obtained from the SPIROMICS (52) cohort through NHLBI BioLINCC (https://biolincc.nhlbi.nih.gov/studies/).

### Statistical analysis

For statistical analysis, Student’s t test was used for comparisons between two groups and comparisons with 3 or more groups were analyzed using One-way ANOVA. For all experiments, *P* value < 0.05 was considered statistically significant. Within individual figure panels *P*-value information is indicated as follows: * = *P* < 0.05, ** = *P* < 0.01, *** = *P* < 0.001 with symbols superimposed (and comparison groups indicated) on graphs accordingly. Correlation analysis was performed based on Pearson’s coefficient. For all data in which 3 or more independent measurements are reported, data are displayed as mean ± standard error of mean (SEM).

### Study Approvals

All human studies were conducted with protocols approved by the Washington University Institutional Review Board and written informed consent was obtained from study participants.

### Recombinant IL-33 protein expression and purification

Splice variants of IL-33 protein were cloned as per previous (12) into pCDH lentiviral vectors or into pHLSEC (53) for expression in Expi293 system (Invitrogen). For mammalian expression, Flag-IL-33^Δ34^-His in pHLSEC was transfected into Expi293 cells using Hype293 (OZ Biosciences) and protein was purified by tandem anti-Flag (GenScript) and NiNTA resins according to protocol. Recombinant protein and co-purified bands were visualized on SDS-PAGE and western blot performed for confirmation of bands. Co-purified HSP70 and HSC70 were confirmed by Mass Spectrometry (methods per MTAC core, see below). Recombinant IL-33^Δ34^ for binding experiments was cloned into pET21a (Novagen) with an N-terminal hexahistidine (6XHis) tag, followed by an N-terminal AviTag™ sequence (GLNDIFEAQKIEWHE) and (GGS)^2^ linker. The IL-33^Δ34^ construct was co-transformed with BirA ligase for in-situ biotinylation as needed. HSP70 was cloned with an N-terminal hexahistidine (6XHis) tag into pETDuet vector (Novagen). Mutants were generated using the Agilent QuickChange II XL kit per manufacturer protocol. Constructs were transformed into Rosetta2(DE3) *E. coli*, (Novagen) under antibiotic selection and bacteria were grown in Luria Broth (LB) at 37°C to an OD_600_ of ∼0.8 before induction with 0.5mM of IPTG at 20°C overnight.

Protein purification from cell pellets was performed by lysis with sonication in 50 mM Na_2_HPO_4_ pH 8.0, 300mM NaCl, 5mM imidazole, 10mM Δ-ME and 10% glycerol with 50 mg/ml Lysozyme (bacterial produced protein), 1 mg DNAse, 1 mM MgCl_2_, 1mM phenylmethylsulfonylfluoride (PMSF). Following centrifugation at 7,000 x g, supernatants were passed over NiNTA superflow resin (Qiagen) with increasing imidazole concentration according to manufacturer protocols based on buffer: Na_2_HPO_4_ pH 8.0, 300 mM NaCl, 10 mM imidazole, 10% glycerol, 5 mM DTT and 10% glycerol supplemented with protease inhibitor cocktail (Sigma). Proteins further purified by size exclusion chromatography (SEC) using a Superdex™ 75 Increase column for IL-33 and a Superdex™ 200 Increase column for HSP70 purifications.

### IL-33^Δ34^ and HSP70 Co-Immunoprecipitation (Co-IP)

For IL-33^Δ34^ and HSP70 co-precipitation analysis using purified proteins, 5 μg of purified biotinylated IL-33^Δ34^ was bound to Pierce Streptavidin Magnetic Beads for 15 mins at 4°C in PBS 0.05% Tween20. Beads were washed 3x and incubated with 7.5 μg of HSP70 for 30 mins at 4°C followed by 3 washes. Proteins were eluted by boiling in 1x SDS PAGE buffer and analyzed by western blot using Invitrogen iBlot system and anti-HSP70 antibody according to reagent list.

### MiniTurboID Proximity Ligation Assay

MiniTurboID was cloned as an N-terminal fusion to IL-33^Δ34^ with (GGS)^2^ linker on a pCDH lentivirus vector backbone (VectorBuilder). Lentivirus was prepared as previously reported (12) and HBE-1 cells were transduced and selected for stable expression. Proximity biotinylation was performed as previously described (32). Briefly, cells were switched to media containing 500 μM exogenous biotin and incubated for 2h, washed, lysed in 700 μL RIPA lysis buffer and incubated with 30 μL of strep magnetic beads at 4°C for 1h with rocking. Beads were washed 2x with RIPA buffer, 1x with 1 M KCl, 1x with 0.1 M Na_2_CO_3_, 1x with 2 M urea 10 mM Tris-HCl (pH 8.0), and 2x with 1 mL RIPA buffer. For trypsin digest and proteomic analysis, biotinylated proteins were eluted in 1x SDS PAGE loading buffer with 20 mM DTT and 2 mM biotin. Trypsin digest and mass spectrometry were performed according to below.

### Lipid binding ELISA

Lipid binding ELISA was performed according to previous (54). Briefly, Nunc MaxiSorp 96-well plates (Thermo) were coated with 5 μg/mL of phosphatidylserine (PS) or phosphatidylcholine (PC) (Avanti polar lipids) diluted in methanol, allowed to air dry and blocked with PBS 1% BSA for 2h. Plates were washed 3x with PBS 0.5% Tween. Non-biotinylated IL-33^Δ34^ (500 ng/ml) +/- HSP70 (500 ng/mL) were bound at room temp for 30 mins, plates washed 3x and bound IL-33^Δ34^ was detected using R&D DuoSet Human IL-33 biotinylated detection antibody following manufacturer protocols and developed using streptavidin HRP (R&D Systems) and TMB peroxidase substrate (SeraCare).

### Quantitative secretion assays

HBE-1 cells were cultured on collagen coated tissue culture plates in UNC BEGM media and U937 cells cultured in RPMI/10% FBS/pen-strep (unless otherwise indicated). All secretion assays were performed at 37°C and 5% CO_2_. Media was exchanged to fresh pre-warmed media at beginning of assay and plates were incubated for 2h. Supernatant was transferred to anti-IL-33 or HSP coated ELISA assay plate and cells were lysed in MPER (Pierce) supplemented with HALT protease inhibitor (Pierce). Lysates were diluted 1:20 in PBS for HSP70 measurements. For all secretion assays, protein was quantified in supernatant and lysate using R&D commercial kits with total assay protein (supernatant + lysate) and % secretion (supernatant/ (supernatant + lysate) x100) quantified based on standard curve.

For chemical inhibition assays, all chemicals were solubilized in DMSO and filter sterilized prior to use. As a control, DMSO was used at highest concentration required for solubility in respective assay. Inhibitors were pre-incubated with cells for 1 hr prior to beginning secretion assay (GW4869, GGA) or 24 hr (Gefitinib, Pifithrin, Rapamycin, Bafilomycin A) and maintained in media during assay. Chemical concentrations used for inhibition assay are as follows: PBS; DMSO vehicle control (2.5%); GW4869 (20 mM), Geranylgeranylacetone (GGA) (20 uM), Pifithrin (10 uM) and Gefitnib (1 uM), Rapamycin (100 nM), and Bafilomycin A (Baf A, 100 nM).

### Heat shock proteins lentiviral knockdown in HBE-1

HSP70 or HSC70 expression knockdown was carried out using commercially available lentivirus encoding 3-4 shRNA constructs targeting GOI (Santa Cruz) according to previous (12). To summarize, 1e5 IFU of virus was combined with TransDux MAX transduction reagent (SBI) and 1e5 HBE-1 cells in BEGM media and placed under selection after 24h incubation. After recovery from selection, cells were used for experimentation, with an aliquot retained for validation of knockdown by qPCR.

### Bronchoalveolar lavage extracellular vesicle (EV) isolation

Extracellular vesicles were isolated from bronchial wash fluid as per previous and in accordance with MISEV2024 guidelines (12, 55). Briefly, bronchial wash was centrifuged at 3000xg to clarify, concentrated and fractionated on a qEV 35 (Izon Science) size exclusion column. EV analysis was performed using ExoView R100 imaging and transmission electron microscopy (not shown) or cryo-EM imaging. Isolated EV preps were submitted for Mass Spectrometry analysis per below.

### Mass Spectrometry Proteomics

Mass Spectrometry analyses were performed by the Mass Spectrometry Technology Access Center at McDonnell Genome Institute (MTAC@MGI) at Washington University School of Medicine. Briefly, samples were reduced, alkylated, and digested with trypsin according to core facility protocols. Digested peptides were desalted on C18 spin columns and analyzed by mass spectrometry. Data was searched against a Human database using MaxQuant search engine and then performed label-free quantification (LFQ) based on the MS1 peptide intensity.

Extracellular vesicle-derived protein was filtered for >1 unique peptide, which yielded 577 proteins, including several EV marker proteins such as ANXA1, ANXA2, ACTN4, CD81, CD9. Obtained LFQ intensities were Log2 transformed, and data were grouped into groups: MTS15 (asthma), MTS16 (nonsmoker), COPD34 (COPD I-III), and COPD53 (COPD IV). These were filtered to retain proteins with at least 60% intensity values in at least one group which resulted in 565 proteins. Imputation was applied to fill in missing values based on normal distribution, which is recommended for label-free quantification. Finally, t-test was carried out and results visualized by volcano plot to determine significant differences in 6 comparison sets.

MiniTurboID labeled proteins were in-gel digested with trypsin using core facility optimized protocol. Peptides were subject to mass spec analysis and the data searched against a human database. In total, 238 proteins were identified including the presence of several naturally occurring biotin proteins that are used as positive controls for biotinylated enrichment experiments (ACACA, PC, MCCC1, MCCC2, PCCA). Results were visualized using Scaffold 5.0.

### Receptor binding ELISA assays

For ST2 and RAGE binding assays ELISA plates were coated with 1 mg/mL of commercial soluble ST2 diluted in PBS or with 0.5 mg/mL anti-Human-Fc (BD) followed by capture of sRAGE-Fc chimera according to manufacturer instructions. Plates were blocked with PBS 1% BSA for 1h and IL-33^Δ34^ was added at 500 ng/ml with or without HSP70 (500 ng/ml). Proteins were diluted in PBS immediately before assay for reduced conditions and incubated in serum free media at 500 ng/ml with or without HSP70 (500 ng/ml) for 4h at 37°C. IL-33^Δ34^ binding was detected using R&D IL-33 detection antibody and developed using streptavidin HRP (R&D Systems) and TMB peroxidase substrate (SeraCare).

### IL-33 receptor cell surface staining

For surface staining, cells were plated in 8 well chamber slides and allowed to reach ∼70% confluency. IL-33^Δ34^ was diluted in serum free media to a final concentration of 100 ng/ml. For competition conditions, HSP70 was added at a concentration of 300 ng/mL and sRAGE-Fc chimera or sST2 at a final concentration of 500 ng/ml. Cells were incubated for 15 min at 25C, washed and fixed with 4% PFA for 5 min. Cells were then blocked for 1h with 2% BSA/1% Fish Gel in PBS. Surface-bound IL-33 was stained with Nessy-1 (1:1000) and anti-mouse-AF647 (1:1000) for A549 staining or polyclonal rabbit anti-IL-33 (1:1000) and then signal amplified using VectaFluor Excel 594 Amplifier Kit (Vector labs).

### Signaling Assays

HEK-Blue IL-33 reporter cells (Invivogen) were used to assay ST2 signaling, and A549-Dual cells were used to assay RAGE signaling. For signaling 30,000 cells were seeded per well of a 96-well plate. For receptor signaling, proteins were diluted to 100 ng/ml (IL-33^Δ34^) and 300 ng/ml (HSP70) in serum free DMEM or Ham’s F-12 media. Cells were switched to serum free media containing IL-33^Δ34^ +/- HSP70 and incubated at 37°C for 2h. Cells were then switched back to full media overnight and SEAP detection was carried out following manufacturer instructions (Invivogen).

## SUPPLEMENTARY FIGURES AND METHODS

**Figure S1.** Associated with Fig. 2.

**Figure S2.** Associated with Fig. 3.

**Figure S3.** Associated with Fig. 5.

**Table S1 & S2.** Human subject demographics & clinical data.

**Table S3.** List of reagents associated with Methods.

## SUPPLEMENTARY FIGURES

**FIGURE S1.**
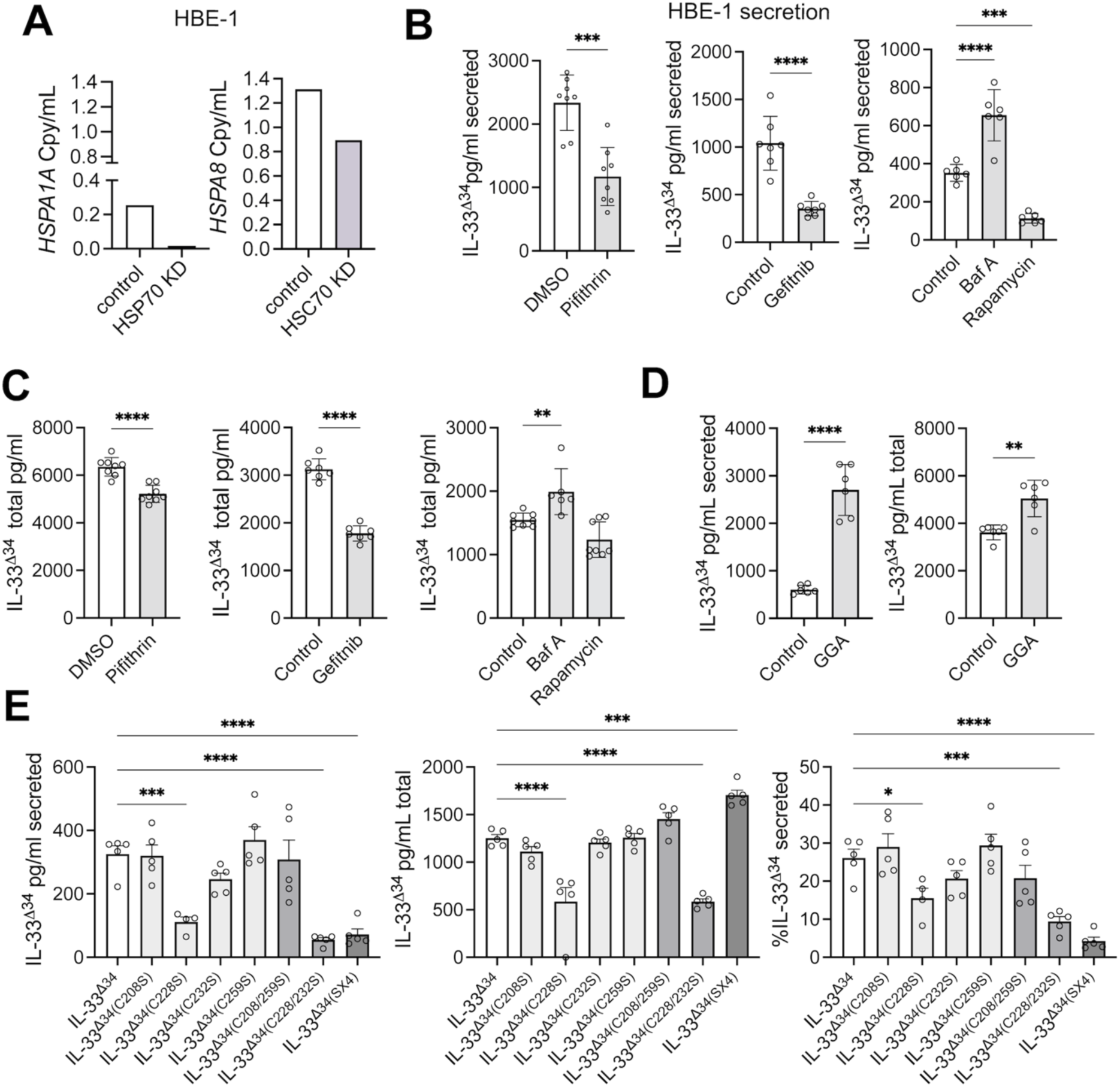
IL-33 expression and secretion assays in HBE-1 cells. A) Validation of shRNA knockdown for HSP70 (*HSPA1A*) and HSC70 (*HSPA8*) in HBE-1 cells by qPCR represented as copies/ml (Cpy/ml) based on plasmid standard. B, C, D) Lentiviral-expressed Flag-IL-33^Δ34^-His secreted (B) and total protein (C) measured by ELISA from HBE cell supernatant under conditions of inhibitors as shown (Pifithrin (PES) 1 μM, Gefitinib 1 μM, Bafilamycin A (Baf A) 100 nM, Rapamycin 100 nM, and (D) GGA 20 μM). E) Flag-IL-33^Δ34^-His secretion data under conditions of single or double Cys to Ser mutants as indicated inclusive of the 4 Cys to Ser mutant (SX4) and wildtype residues C208, C228, C232 and C259. Data points displayed with mean ± SEM. *P*-value limits: \**P* < 0.05, \*\**P* < 0.01, \*\*\**P* < 0.001, \*\*\**P* < 0.0001.

**FIGURE S2.**
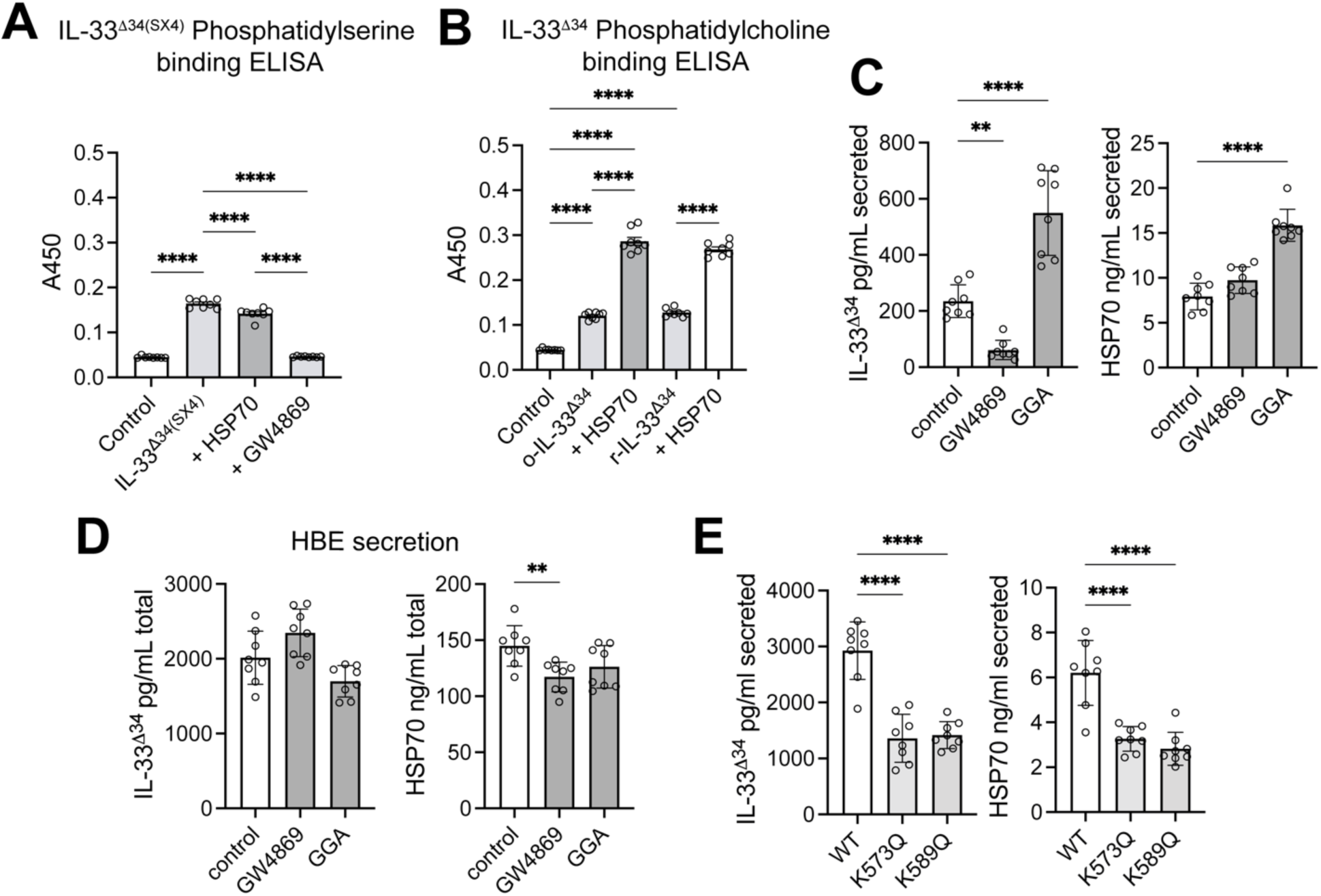
A) Phosphatidylserine (PS) binding for IL-33^Δ34(SX4)^ mutant (C208S, C228S, C232S, C259S) and B) phosphatidylcholine (PC) for IL-33^Δ34^ binding ELISAs under conditions of HSP70 (500 ng/ml) and GW4869 (20 μM) in competition. Detection was performed using biotinylated anti-IL-33 secondary antibody (R&D Systems) and Streptavidin HRP, and TMB substrate followed by A450 absorbance measurements. C, D) Secreted and total Flag-IL-33^Δ34^-His and endogenous HSP70 in HBE-1 cells under conditions of GW4869 (20 μM) and GGA (1 μM) treatment, detection using anti-IL-33 and HSP70 Duosets (R&D Systems) and TMB substrate. E) Data points displayed with mean ± SEM. *P*-value limits: \**P* < 0.05, \*\**P* < 0.01, \*\*\**P* < 0.001.

**FIGURE S3.**
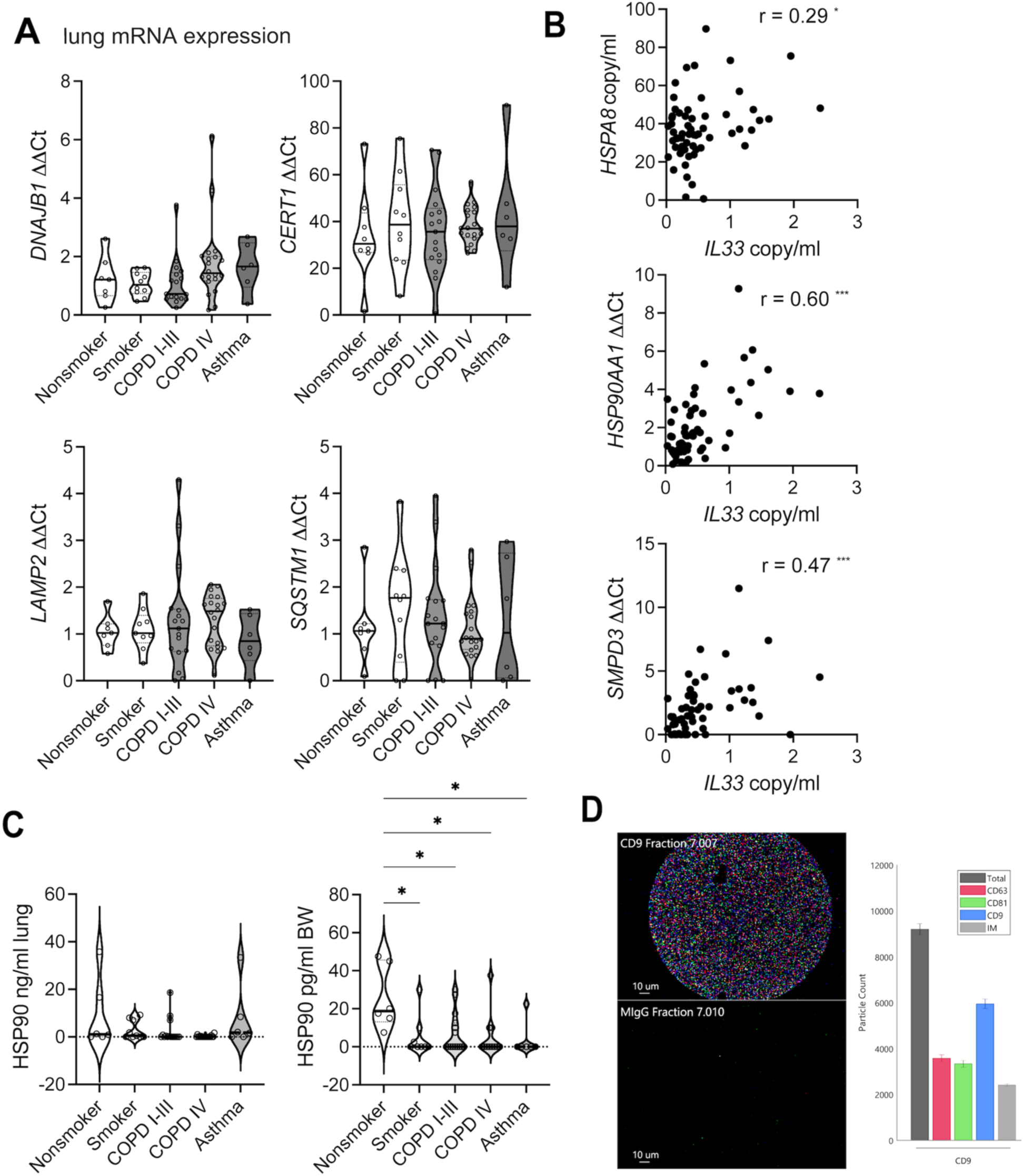
A) Lung tissue qPCR for Heat shock and proteostasis intermediates (HSP40/*DNAJB1*, p62/*SQSTM1*) and vesicular trafficking intermediates (*LAMP2*, *CERT1*) in nonsmokers (n=7), smokers (n=11) COPD I-III (n=17), COPD IV (n=20) and severe asthma (n=6) lung tissue specimens represented as fold-change by 1′1′Ct method and normalized to GAPDH. B) Pearson’s correlation for IL33 expression with HSC70/*HSPA8*, HSP90/*HSP90AA1* and nSMase2/*SMPD3*. C) HSP90 protein levels measured in lung tissue specimens and bronchial wash fluid from representative specimens in A) including nonsmokers (n=6), smokers (n=8) COPD I-III (n=16), COPD IV (n=20) and severe asthma (n=6). D) Tetraspanin (CD63, CD81 and CD9) positive vesicles were verified using Exoview R100 analysis platform following manufacturer protocols. A representative image of CD9 and negative control capture spots are represented. Particle counts are averaged from 3 different CD9 capture spots. Data points displayed as mean ± SEM. *P*-value limits: \**P* < 0.05.

**Table S1.** Demographic and clinical data for patient specimens in Washington University cohort: Lung transplant donor specimens without COPD (Nonsmoker), rejected donors without COPD (smoker) or mild-moderate (GOLD Stage I-III) COPD^#^ and lung transplant recipients with severe (GOLD Stage IV) COPD. ^#^Rejected donor lungs grouped as COPD I-III separately from smokers based on the following criteria: 1) presence of emphysema on chest CT, 2) hyperinflation/air trapping on gross pathological examination and 3) positive tobacco history >10 pk-yr. *Severe asthma clinical history with cause of death status asthmaticus.

| Characteristics | Nonsmoker | Smoker | COPD I-III<sup>#</sup> | COPD IV | Asthma<sup>*</sup> |
| --- | --- | --- | --- | --- | --- |
| Number per group | 7 | 11 | 17 | 20 | 8 |
| Mean age (range) | 17.7 (3-32) | 40.8 (21-67) | 51.9 (12-71) | 61.4 (52-70) | 35 (24-44) |
| Male:Female:Unknown | 4:0:3 | 3:2:6 | 3:1:10 | 7:13 | 2:2:2 |
| FVC (L) | Unknown | Unknown | Unknown | 1.9655 | Unknown |
| FEV1 (L) | Unknown | Unknown | Unknown | 0.5235 | Unknown |
| FEV<sub>1</sub>/FVC (%) | Unknown | Unknown | Unknown | 24.95 | Unknown |
| Pack-years (range) | N/A | Unknown | 40.8 (30-60) | 42.7 (10-96) | Unknown |

**Table S2.** Clinical characteristics of COPD patients based on GOLD Stage obtained from SPIROMICS cohort (52).

| Characteristics | COPD 0 | COPD 1 | COPD 2 | COPD 3 |
| --- | --- | --- | --- | --- |
| Number per group | 91 | 28 | 29 | 4 |
| Mean age (range) | 56.6 (40-74) | 64 (46-75) | 62.8 (52-74) | 53.5 (41-68) |
| Male:Female | 40:51 | 16:12 | 18:11 | 3:1 |
| FVC (L) | ND | ND | ND | ND |
| FEV1 (L) | 4 | 2.27 | 1.53 | 1.84 |
| FEV<sub>1</sub>/FVC (%) | ND | ND | ND | ND |
| Pack-years (range) | 30.9 (0-112) | 51.9 (20-118) | 58.4 (25.5-108) | 58.9 (25-90) |

**Table S3:** List of reagents and materials associated with STAR Methods.

| Reagent | Supplier | Reference/Identifier |
| --- | --- | --- |
| **Antibodies and Proteins** |  |  |
| Human IL-33 rabbit polyclonal (1:50 IF) | Sigma Aldrich | HPA024426 |
| Human IL-33 goat monoclonal (1:1000 WB) | R&D systems | MAB36253 |
| Human IL-33 mouse monoclonal (Nessy-1) (1:1000 IF) | Enzo | ALX-804-840 |
| Human HSP70 antibody (6B3) (1:5000 WB, 1:100 IF) | Cell Signaling | 4873S |
| Human HSP90 antibody (1:100 IF) | Proteintech | 60318-1-1g |
| Soluble human RAGE-Fc | R&D systems | 1145-RG |
| Soluble human ST2 | R&D systems | 11272-ST |
| Anti-human Fc | BD | 32935S |
| Anti-rat-HRP | Santa Cruz | sc-2032 |
| Anti-rabbit vectafluor 594 kit | Vector labs | DK-1594 |
| Anti-rat-488 | Invitrogen | A21208 |
| IF = immunofluorescence, WB = western blot |  |  |
| **Commercial lentiviruses** |  |  |
| Scrambled shRNA | Santa Cruz | sc-108080-V |
| HSC70 shRNA | Santa Cruz | sc-62655-V |
| HSP70 shRNA | Santa Cruz | sc-29352-V |
| **Biological Samples** |  |  |
| SPIROMICS BAL | NHLBI | N/A |
| Control and COPD lung tissue | BJH, UNMC | N/A |
| Control and COPD Bronchial wash fluid | BJH | N/A |
| **Chemicals, Peptides, and Recombinant Proteins** |  |  |
| GW4869 | Sigma Millipore | D1692 |
| Phosphatidylserine | Avanti Polar lipids | 840032P |
| Phosphatidylcholine | Sigma Aldrich | 63556 |
| Pifithrin | Sigma Aldrich | P0122 |
| Gefitinib | Sigma Aldrich | SML1657 |
| Bafilomycin A | Sigma Millipore | B1793-10UG |
| Rapamycin | Sigma Aldrich | R8781 |
| Geranylgeranylacetone | Sigma Aldrich | G5048 |

| Commercial Assays |  |  |
| --- | --- | --- |
| IL-33 ELISA Duoset | R&D Systems | DY3625 |
| HSP70 ELISA Duoset | R&D Systems | DY1663 |
| HSP90 ELISA | Proteintech | KE00054 |
| Cell Lines |  |  |
| HBE-1 | Marisco Lung Institute |  |
| HEK-Blue | Invivogen |  |
| A549 Dual | Invivogen |  |
| U937 | ATCC |  |
| qPCR assays | IDT PrimeTime® predesigned qPCR assays |  |
| HSPA1A | 421177696 |  |
| HSPA8 | 421177700 |  |
| IL33 | 441084120 |  |
| STIP1 | 454622465 |  |
| STUB1 | 454622457 |  |
| HSP90AA1 | 416230057 |  |
| SMPD3 | 218154457 |  |
| MUC5AC | 166039047 |  |
| DNAJB1 | 424729991 |  |
| CERT1 | 414584019 |  |
| LAMP2 | 206830890 |  |
| SQSTM1 | 454622461 |  |
| CCT5 | 508892978 |  |
| CCT7 | 508892982 |  |
| Constructs |  |  |
| human Flag-IL-33<sup>Δ34</sup>-His<br>C208S<br>C228S<br>C232S<br>C259S<br>C208/259S<br>C228/232S<br>C208/228/232/259S | pCDH – puro, neo |  |
| human IL-33<sup>Δ34</sup>-birA-6His | pET21 |  |
| human IL-33<sup>Δ34</sup>-MiniTurbo-V5 | pCDH - neo |  |
| Human HSP70-His | pCDH - neo |  |
| Human HSP70 (K573Q)-His | pCDH - neo |  |
| Human HSP70 (K589Q)-His | pCDH - neo |  |

